# Hexokinase 1-mediated plant immunity under abiotic stress depends on chloroplast positioning

**DOI:** 10.1101/2024.12.04.626761

**Authors:** Sophia Bianca Bagshaw, Anastasia Kitashova, Beyza Özmen, Chun Kwan Yip, Bianca Emily Süling, Laura Schröder, Tatjana Kleine, Thomas Nägele

**Affiliations:** LMU München, Faculty of Biology, Plant Evolutionary Cell Biology, Großhaderner Str. 2-4, 82152 Planegg, Germany; LMU München, Faculty of Biology, Plant Development, Großhaderner Str. 2-4, 82152 Planegg, Germany; LMU München, Faculty of Biology, Plant Molecular Biology, Großhaderner Str. 2-4, 82152 Planegg, Germany

**Keywords:** *Arabidopsis thaliana*, cold acclimation, chloroplast positioning, carbohydrate metabolism, plant immune response

## Abstract

Molecular networks involved in the responses of plants towards environmental changes are multifaceted and affect diverse metabolic and signalling pathways. Under challenging environmental conditions, such as low temperatures and high light intensities, plants need to immediately adjust their metabolism to prevent irreversible tissue damage, e.g., caused by reactive oxygen species. Regulation of photosynthesis and carbohydrate metabolism plays a crucial role in this stress response. Here, we analysed mutants of *Arabidopsis thaliana*, which were affected in either central enzymatic activities of carbohydrate metabolism, in chloroplast positioning or in a combination of both. Plants were exposed to a treatment of combined cold and elevated light. While mutants with deficiencies in sucrose (*spsa1*) or starch (*pgm1*) metabolism showed affected metabolic pathway regulation under abiotic stress, Hexokinase 1 mutants (*hxk1*) showed a severe growth phenotype with lesions and pale areas on leaf tissue. In a double mutant, which combined deficiencies in Chloroplast Unusual Positioning 1 (CHUP1)-mediated chloroplast positioning and HXK1 (*chup1 x hxk1*), this growth phenotype vanished resulting in wild type-like plants. Transcriptome analysis revealed a significantly affected immune response of *hxk1* plants, which was suppressed in the double mutants. Our findings provide evidence for a role of HXK1 as a positive regulator of the plant immune response. Finally, we suggest that, due to its potential role as a negative regulator of plant immunity, CHUP1 deficiency counteracted the reduced immunity of *hxk1* in the double mutant which rescued the plants. Future studies might now reveal whether deficiencies in CHUP1 function and/or transcription represent a conserved strategy to increase plant immunity under abiotic stress.

## Introduction

Plant carbohydrates are direct products of photosynthetic CO_2_ assimilation. Within the Calvin-Benson-Bassham cycle (CBBc), ribulose-1,5-bisphosphate carboxylase/ oxygenase (Rubisco) enzymes catalyse both carboxylation and oxygenation reactions resulting in two or one molecule(s) of 3-phosphoglycerate, respectively. In the following reactions, which are ATP-and NADPH-dependent, triose phosphates are synthesised, which are substrates for either metabolic pathways or regeneration of CBBc intermediates. Both ATP and NADPH are synthesised within the primary photosynthetic reactions across thylakoid membranes in chloroplasts (Govindjee et al., 2017). To prevent an overaccumulation of ATP/NADPH and a depletion of ADP and NADP^+^, a tight regulation of the photosynthetic electron transport and ATP/NADPH consumption in the CBBc is necessary. One way the activity of enzymes central to the CBBc can be regulated in response to the photosynthetic electron transport rate is via thioredoxins which, in a ferredoxin-dependent manner, affect the reduction state and the activity of their targets (Geigenberger et al., 2017). The carbohydrate metabolism in the chloroplast is also tightly connected with cytosolic pathways through mechanisms such as metabolite transport across the chloroplast envelope (Flügge, 1999; Patzke et al., 2019). An example of this is the translocation of plastidial triose phosphates in an antiport manner into the cytosol while inorganic phosphate is transported into the chloroplast which prevents phosphate depletion and deregulation of photosynthetic CO_2_ assimilation (Flügge and Heldt, 1981). In the cytosol, sugar phosphates are substrate for energy metabolism, numerous metabolic pathways, sucrose biosynthesis, and biomass accumulation. Together with starch, sucrose represents the central product of photosynthesis and is, in many plant species, the most abundant transport sugar which supplies non-photosynthetic and below-ground leaf tissue with carbon equivalents (Braun, 2022).

Due to its central role in whole plant metabolism, stress tolerance and developmental processes, understanding the regulation of sucrose metabolism in a changing environment is important for many research areas. However, the experimental analysis of sucrose metabolism is aggravated by the diversity of factors that influence actual sucrose concentrations. The cyclic biosynthesis and breakdown of sucrose, catalysed by sucrose phosphate synthase, invertases, and hexokinases, challenges the interpretation of concentration dynamics (Nägele, 2022). Such dynamics are typically observed if environmental factors, like temperature or light intensity, change. Under low temperatures, sucrose concentrations significantly increase together with many other carbohydrates, amino acids and polyamines (Hannah et al., 2006; Guy et al., 2008; Hoermiller et al., 2022). These rising concentrations have been discussed in context of cryoprotection of membranes, where polar head groups, e.g., of carbohydrates and membrane lipids, are suggested to interact and prevent membrane fusion (Hincha et al., 2003). Further, chloroplast-located raffinose family oligosaccharides (RFOs) were shown to stabilize Photosystem II during freeze/thaw cycles, most probably due to stabilization of thylakoid membranes and/or membrane-localized proteins (Knaupp et al., 2011). Sucrose, together with galactinol, is substrate for RFO biosynthesis which again emphasises its multifaceted metabolic role in plant metabolism.

By relocating chloroplasts within a cell, plants can regulate electromagnetic energy absorption. Under high light intensities, in an avoidance response, chloroplasts are positioned along the anticlinal cell wall to reduce the total surface area of light absorption (Kasahara et al., 2002). In contrast, under low light intensities, chloroplasts accumulate perpendicular to the incident light to maximise absorption (accumulation response). The accumulation response was found to be mediated by blue light receptors Phototropin 1 and 2 (PHOT1 and 2; (Sakai et al., 2001)), while the avoidance response is mediated by PHOT2, which was also observed to affect chloroplast localization under low temperature (Kagawa et al., 2001; Kodama et al., 2008). The organelle movement itself is catalysed by Chloroplast Unusual Positioning 1 (CHUP1) proteins, which represent plant-specific actin polymerization factors (Kong et al., 2024). Recent work suggested that chloroplast actin filaments, which are built at the interface between plasma membrane and chloroplasts, generate the motive force during chloroplast movement but detailed mechanisms remain to be elucidated (Wada and Kong, 2018; Kong et al., 2024).

Due to the observations that (i) chloroplast positioning significantly affects photosynthetic performance, (ii) photosynthetic performance affects carbohydrate metabolism, and (iii) carbohydrate metabolism plays an essential role for acclimation to a changing environment, we analysed mutants of *Arabidopsis thaliana* which were either affected in enzyme activities of the central carbohydrate metabolism, or in chloroplast positioning, or in a combination of both. Applying a combined elevated light and low temperature treatment revealed that Hexokinase 1 (HXK1) plays a central role for growth under such growth conditions which was linked to chloroplast positioning and plant immune response.

## Materials and methods

### Plant material and growth conditions

Plants of wild type *Arabidopsis thaliana*, accession Col-0, and seven mutants deficient in the chloroplast movement protein CHLOROPLAST UNUSUAL POSITIONING 1 (*chup1*, AT3G25690, SALK_129128C), plastidial phosphoglucomutase 1 (*pgm1*, AT5G51820, CS3092), cytosolic sucrose phosphate synthase A1 (*spsa1*, AT5G20280, SALK_148643C), cytosolic hexokinase 1 (*hxk1*, AT4G29130, SALK_034233C), as well as double mutants *pgm1 x chup1 (pc)*, *spsa1 x chup1 (sc)*, and *hxk1 x chup1 (hc)*, were grown on a 1:1 mixture of GS90 soil and vermiculite in a climate chamber under controlled short day conditions (8 h/16 h light/dark; 100 μmol m^−2^ s^−1^; 22°C/16°C; 60% relative humidity). After two weeks, plants were transferred to a greenhouse and grown under long day conditions (16 h/8 h light/dark; 100 μmol m^−2^ s^−1^; 22°C/16°C; 60% relative humidity). After one further week, seedlings were either (i) sampled at midday, i.e., after 8 h of light (0 days of acclimation), or (ii) transferred to a cold room for low temperature and elevated light treatment (LT/ET; 16 h/8 h light/ dark; 250 μmol m^−2^ s^−1^; 4°C/4°C). LT/ET exposed plants were sampled after 4 days at midday, i.e., after 8 h of light. Each sample consisted of four pots with densely grown seedlings (Figure 1), which were immediately frozen in liquid nitrogen, ground to a fine powder, and lyophilised. All mutations were verified by PCR. Mutations of PGM1, SPS and HXK1 were also validated by quantifying starch amounts as well as SPS and glucokinase activities, respectively (Supplementary Table ST1). Mutation of CHUP1 was additionally validated by microscopy (Supplementary Figure SF1).

**Figure 1.**
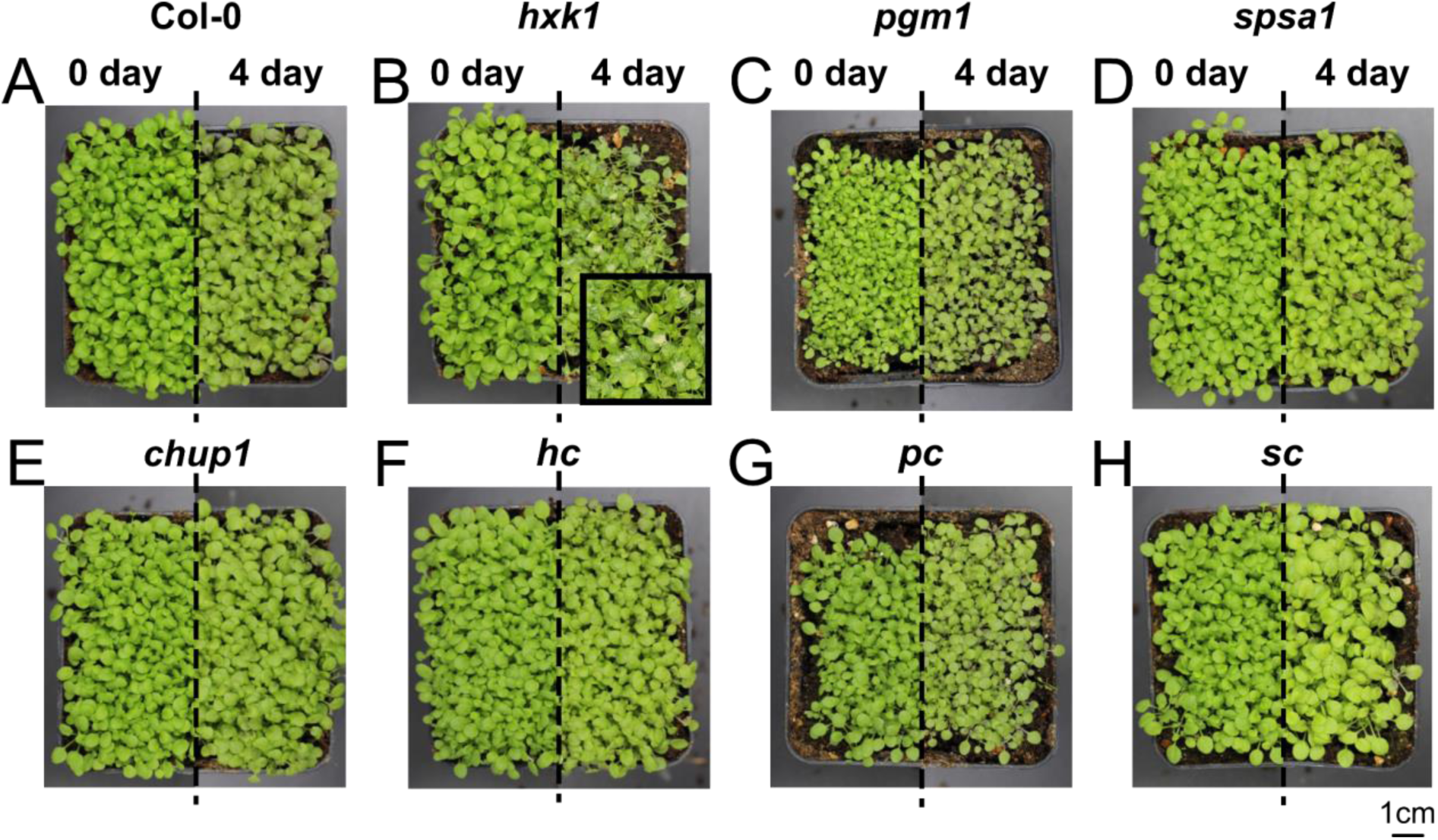
Effects of deficiencies in enzymatic activities on growth and stress response. Each genotype is shown before (0 day) and after (4 day) exposure to combined cold/high light treatment (left/right). **(A)** Col-0, **(B)** *hxk1*, **(C)** *pgm1,* **(D)** *spsa1,* **(E)** *chup1,* **(F)** *hc,* **(G)** *pc*, **(H)** *sc.* Magnification in **(B)** shows the *hxk1* phenotype under LT/EL.

### Chlorophyll fluorescence measurements

Chlorophyll fluorescence parameters were recorded at ambient temperature (22°C) with Imaging-PAM MAXI version (Heinz Walz GmbH; www.walz.com). Maximum quantum yield of PSII (Fv/Fm) was determined after 15 min of dark adaptation by supplying a saturating light pulse. Dynamics of quantum efficiency of PSII (Y(II)), electron transport rate (ETR), photochemical (qP), non-photochemical quenching (qN, NPQ), quantum yield of regulated energy dissipation (Y(NPQ)) and quantum yield of nonregulated energy dissipation (Y(NO)) were determined after 15 min of light adaption to either (i) 100 μmol m^−2^ s^−1^ for non-acclimated plants; or (ii) 250 μmol m^−2^ s^−1^ for LT/EL-acclimated plants.

### Pigment extraction and quantification

Chlorophyll was extracted by solubilising dried plant material in 80% acetone before samples were incubated on ice for 10 min. Following a short centrifugation at 4°C, absorbance at 652 nm was determined in the supernatant and chlorophyll content was calculated as described earlier (Bruinsma, 1961).

Anthocyanins were quantified photometrically as described before (Kitashova et al., 2023). Lyophilized plant material was incubated with 1M HCl at 25° C for 30 min. The samples were then centrifuged at 20,000g, and the supernatants were transferred to a new tube. A second extraction was performed at 80° C. After centrifugation, the two supernatants were pooled, and the absorbance was measured at 540 nm. Total anthocyanin concentration was normalised to pelargonidin (C15) standards.

### Quantification of protein amounts

Total protein amount was determined photometrically as described earlier with slight modifications (Fürtauer et al., 2018). In brief, lyophilised plant material was solubilised in 8.8 M urea and 50 mM HEPES, pH 7.8. After filtering out plant material debris, proteins were precipitated using 100% ice cold acetone with 2 mM DTT. Pellets were washed twice with methanol and acetone, and solubilised in 8.8 M urea, 50 mM HEPES, pH 7.8. Protein amounts were quantified using the Bradford solution for protein determination (PanReac AppliChem, www.itwreagents.com).

### Quantification of starch, soluble carbohydrates and hexose phosphates

Transitory starch, soluble carbohydrates, and hexose phosphate amounts were determined as described before (Kitashova et al. 2023). Dried plant material was incubated with 80% ethanol at 80°C for 30 minutes. Following brief centrifugation, the supernatant containing soluble sugars was transferred to a new tube. Starch-containing pellets were incubated with 0.5 N NaOH at 95°C for 60 min before acidification with 1 M CH_3_COOH, and then digested with amyloglucosidase, yielding glucose moieties. Glucose concentration was quantified photometrically in a glucose oxidase/peroxidase/dianisidine reaction at 540 nm.

Soluble sugar-containing ethanol extracts were dried and resolubilised in water. After incubation of samples with 30% KOH at 95°C, sucrose amount was quantified following an incubation with 0.14% (w/v) anthrone in 14.6 M H_2_SO_4_ at 40°C for 30 min, with absorbance detection at 620 nm. Glucose was quantified using a coupled hexokinase/glucose-6-phosphate dehydrogenase (G6PDH) assay, and fructose using a coupled hexokinase/phosphoglucose isomerase/G6PDH assay, which yielded NADPH detectable at 340 nm.

Glucose 6-phosphate (G6P) and fructose 6-phosphate (F6P) were extracted using trichloroacetic acid (TCA) in diethyl ether (16% w/v). Following washing with 16% (w/v) TCA in 5 mM EGTA, samples were neutralised with 5 M KOH/1 M triethanolamine. Samples were incubated at 30°C for 30 min in a buffer containing G6PDH in 1 M Tricin pH 9, 10 mM NADP^+^ (and PGI for quantifying F6P), resulting in the production of NADPH. After the remaining NADP^+^ was depleted by adding 0.5 M NaOH and samples were neutralised with 0.5 M HCl, G6P and F6P concentrations were determined in a cyclic reaction. Samples were supplied with G6PDH in the 1 M Tricin pH 9, 100 mM MgCl_2_, 100 mM EDTA, 100 mM G6P buffer, and mixed with thiazolyl blue and phenazine methosulfate. The produced formazan dye was then detected at 570 nm.

### Quantification of enzyme activities

Activities of sucrose phosphate synthase (SPS), glucokinase (GLCK), fructokinase (FRCK), and invertase (INV) were determined under substrate saturation (v_max_) as described before (Kitashova et al., 2023). For quantification of SPS maximal activity, lyophilised plant material was suspended in 50 mM HEPES pH 7.5, 10 mM MgCl_2_, 1 mM EDTA, 2.5 mM DTT, 10% (v/v) glycerine and 0.1% (v/v) Triton X-100. Following centrifugation at 4°C with 20.000 g, the supernatant was incubated for 30 min at 25°C with 50 mM HEPES pH 7.5, 15 mM MgCl_2_, 2.5 mM DTT, 35 mM UDP-glucose, 35 mM F6P and 140 mM G6P. After the reaction was stopped by boiling samples with 30% KOH, the produced sucrose amounts were determined as described above.

For quantification of glucokinase (GLCK) and fructokinase (FRCK) activities, dried plant material was incubated with 50 mM Tris pH 8.0, 0.5 mM MgCl_2_, 1 mM EDTA, 1 mM DTT and 1% (v/v) Triton X-100, then centrifuged at 4°C at 20.000 g. The supernatant was mixed with 100 mM HEPES pH 7.5, 10 mM MgCl_2_, 2 mM ATP, 1 mM NADP_+_, 0.5 U G6PDH and either 5 mM glucose for glucokinase measurement or 5 mM fructose for fructokinase measurement, and absorbance was measured at 30°C at 340 nm.

Cytosolic (cINV) and vacuolar (vINV) invertase activities were determined following extraction on ice in a buffer containing 50 mM HEPES–KOH pH 7.5, 5 mM MgCl_2_, 2 mM EDTA, 1 mM phenylmethylsulfonylfluoride, 1 mM DTT, 10% (v/v) glycerine and 0.1% (v/v) Triton X-100. Activity of cINV was determined using a reaction buffer with pH 7.5 (20 mM HEPES–KOH, 100 mM sucrose), and vINV using a reaction buffer with pH 4.7 (20 mM sodium acetate, 100 mM sucrose). After incubation of samples at 30°C and stopping the reaction by heating samples to 95°C, glucose moieties were photometrically determined with a coupled glucose oxidase/peroxidase/dianisidine reaction at 540 nm.

### Subcellular quantification of hexose phosphates amounts

G6P and F6P concentrations in plastids and cytosol were determined by combining non-aqueous fractionation (NAF) and photometrical determination of hexose phosphates. A benchtop protocol was applied for NAF (Fürtauer et al., 2016) and adapted as described previously (Kitashova et al., 2024). In summary, lyophilised plant material was homogenised in tetrachloroethylene (ρ = 1.60 g cm^-3^) with an ultrasonic homogenizer (Hielscher Ultrasonics UP200St, www.hielscher.com). Following centrifugation at 20,000 g, the supernatant and pellet were separated, and the density of the supernatant was decreased using heptane (ρ = 0.68 g cm^-3^). The supernatant was sonicated again, and after centrifugation, the pellet and supernatant were separated again. The procedure was repeated for, in total, 6 fractions with densities between 1.35 and 1.60 g cm^-3^. The pellets were resuspended, split into 2 sub-fractions, and dried in a desiccator. These samples were used for (i) marker enzyme quantification; and (ii) hexose phosphate quantification. Activities of the marker enzymes alkaline pyrophosphatase (plastidial) and uridine 5′-diphosphoglucose pyrophosphorylase (cytosolic), were determined photometrically (Fürtauer et al., 2016). Subcellular hexose phosphate amounts were quantified as described above and correlated with marker enzyme activities to reveal an estimate of plastidial and cytosolic F6P and G6P proportions.

### Transmission electron microscopy (TEM)

Seedlings were covered with aluminium foil at the end of the night prior to chemical fixation to minimize starch accumulation. Seedlings were then cut into 1 x 1 mm pieces and fixed with a buffer containing 2.5 % glutaraldehyde, 75 mM cacodylate, 2 mM MgCl_2_, pH 7.0. The fixation schedule further contained washing steps with 1% osmium tetroxide and dehydration steps with en-bloc contrasting containing 1 % uranyl acetate. For embedding, leaf pieces were polymerized in Spurr’s resin of medium rigidity over night at 63 °C. Ultrathin sections of 70 nm were post stained for 2 min with lead citrate and TEM was performed with the Zeiss EM 912, outfitted with an omega filter and lanthanum hexaboride source. The Zeiss EM 912 operated at 80 kV in zero-loss mode. Images were acquired using a 2k x 2k slow-scan CCD camera (Tröndle Restlichtverstärkersysteme, Moorenweis, Germany).

### RNA sequencing and data analysis

Total RNA from plants was isolated using Trizol (Invitrogen, Carlsbad, USA) and purified using Direct-zol™ RNA MiniPrep Plus columns (Zymo Research, Irvine, USA) according to the manufacturer’s instructions. RNA integrity and quality was assessed with an Agilent 2100 Bioanalyzer (Agilent, Santa Clara, USA). Depletion of ribosomal RNA, generation of directional lncRNA-Seq libraries and 150-bp paired-end sequencing (depth of approximately 6G) on an Illumina Novaseq 6000 system (Illumina, San Diego, USA) were conducted at Biomarker Technologies (BMK) GmbH (Münster, Germany) with standard Illumina protocols. Three independent biological replicates were used per genotype.

RNA-Seq reads were analyzed on the Galaxy platform (Jalili et al., 2020) essentially as described (Tang et al., 2024) with one exception: DESeq2 (Love et al., 2014) with the fit type set to “parametric”, “multi-mapping allowed”, a linear two-fold change cutoff, and an adjusted *P* < 0.05, were used to determine the differential expression of both plastid-and nuclear-encoded genes.

### Data analysis and statistics

Data was evaluated and statistically analysed using RStudio (Posit Team, 2024) and MATLAB® (www.mathworks.com). Prior to analysis of variance, statistical outliers of each variable were replaced by medians or mean values. If sample sizes varied within one data table, e.g., between hexose phosphates (n=6) and soluble sugars/enzyme activities (n=5), missing values were filled by mean values. GO term enrichment analysis was performed using PlantRegMap with a threshold p-value ≤ 0.001 (Tian et al., 2020).

## Results

### Seedlings of *hxk1* are affected in growth and leaf metabolism under combined cold and elevated light

Exposure to a combination of low temperature and elevated light intensities (LT/EL) for four days resulted in a clearly distinguishable growth phenotype in seedlings of *hxk1* (Figure 1). Pale leaf areas were only observed in *hxk1* (Fig. 1 B). The *pgm1* and *pc* mutants were the smallest among all genotypes (Fig. 1 C, G). Interestingly, the phenotype of *hxk1* plants was suppressed by the additional mutation of *chup1* in the *hc* double mutant (Fig. 1 F). The growth experiment was repeated, and all phenotypes could be reproduced under these conditions.

The treatment of LT/EL significantly reduced the ratio of fresh weight (FW) to dry weight (DW) in all genotypes (Fig. 2 A). The highest median was observed in the wild type under control conditions (0d) which showed a more than 15-fold higher FW than DW. Lowest medians at 0d were observed for *hxk1* and *sc* which was an approx. 12.5-fold higher FW than DW. After 4d of LT/EL, median ratios dropped below a ratio of 10 in all genotypes except for *pgm1* and *pc,* which showed a ratio of ∼12 and ∼10.5. The lowest ratio was observed for *hxk1* at LT/EL (median FW/DW ∼5).

**Figure 2.**
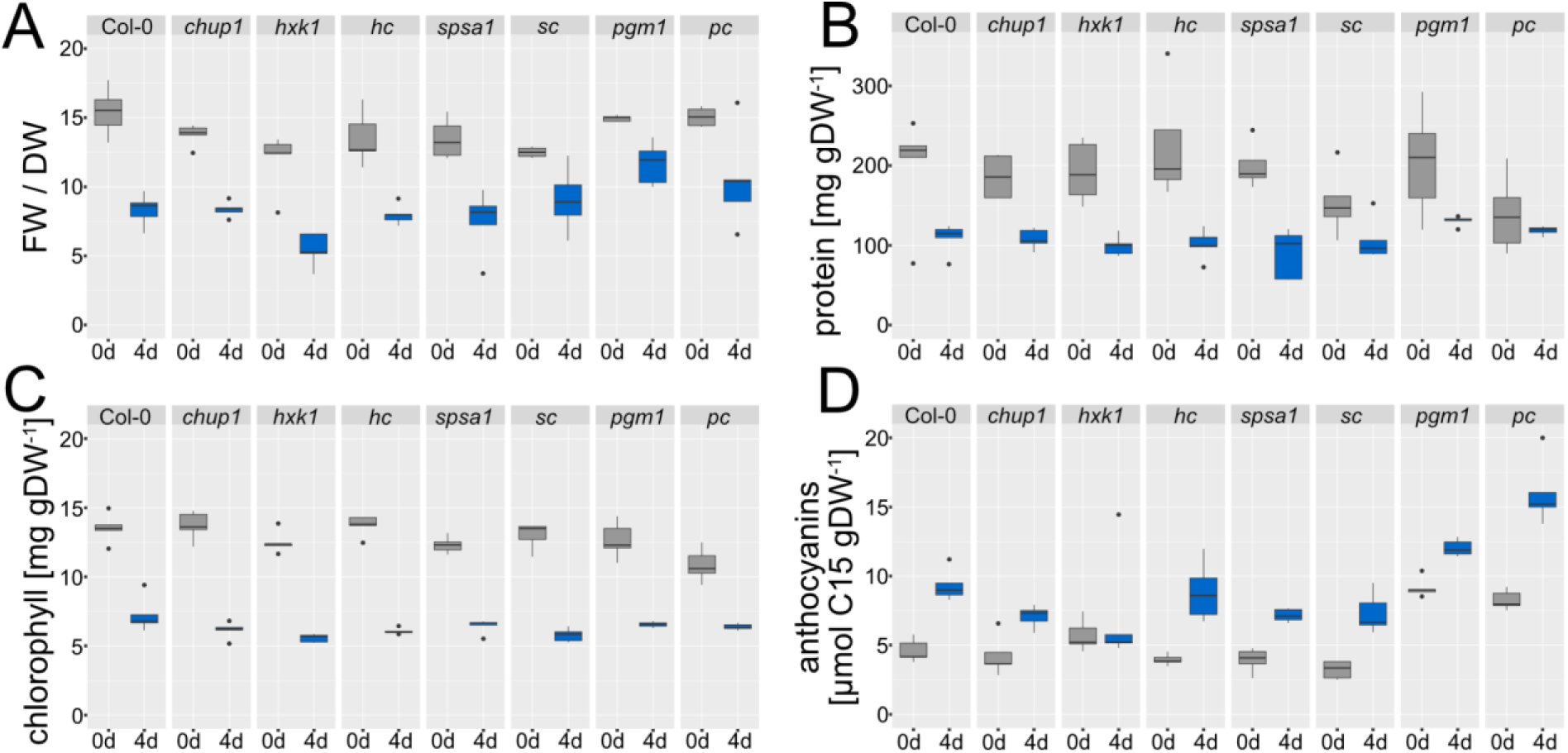
Effects of stress treatment on biomass, proteins and pigment content. **(A)** Ratios of fresh weight (FW) to dry weight (DW). Treatment (LT/EL) had a significant effect in all genotypes (ANOVA, p<0.05). **(B)** Total protein amounts normalised to dry weight. Treatment (LT/EL) had a significant effect in all genotypes except for *pgm1* and *pc* (ANOVA, p<0.01). **(C)** Chlorophyll content normalised to dry weight. Treatment (LT/EL) had a significant effect in all genotypes (ANOVA, p<0.001). **(D)** Anthocyanin content normalised to a C15 standard (pelargonidin) and to dry weight. Treatment (LT/EL) resulted in a significant increase of anthocyanins across all genotypes except for hxk1 (ANOVA, p<0.001). n = 5, where each sample consisted of four pots. Grey boxes: control, i.e., 0d of LT/EL. Blue boxes: treatment, i.e., 4d of LT/EL. The full data set is provided in the supplements (Supplementary Table ST1).

Across all genotypes, the total protein amount at 0d showed a similar median value of ∼200 mg gDW^-1^ except for double mutants *sc* and *pc,* which had a lower content of ∼150 mg gDW^-1^ (Fig. 2 B). After 4d LT/EL, protein amount dropped to ∼100 mg gDW^-1^ and was highest in *pgm1* (∼130 mg gDW^-1^). The double mutant *pc* was the only genotype which did not show a significant reduction of protein amount under LT/EL.

Chlorophyll content at 0d was between 10 and 14 mg gDW^-1^ and significantly dropped in all genotypes to values between 5 and 7 mg gDW^-1^ after 4d LT/EL (Fig. 2 C). Both mutants of *hxk1* and *sc* had a significantly lower chlorophyll content than Col-0 under LT/EL (ANOVA, p < 0.01).

In all genotypes, photometrically determined anthocyanin contents increased significantly after 4d LT/EL except for *hxk1,* which showed a similar median value before and after treatment (Fig. 2 D). Highest amounts of anthocyanins were determined in *pgm1* and *pc* which showed approx. 1.5x-fold higher values than Col-0 under both conditions.

Carbohydrates were strongly and significantly affected in their amounts due to the LT/EL treatment (Fig. 3). Compared to Col-0, the *spsa1* mutant had significantly elevated starch amounts under control conditions (ANOVA, p < 0.01) which was not observed at LT/EL (Fig. 3 A). The *hxk1* mutant was found to accumulate significantly less starch than Col-0, while in the *hc* (and also *sc*) double mutant starch accumulated to similar amounts as in the *chup1* mutant. Treatment-induced sucrose dynamics were significantly affected in both *spsa1* and *sc* which showed the lowest sucrose amounts after 4d LT/EL (Fig. 3 B). In *pgm1* and *pc* mutants, sucrose amount was significantly higher than in Col-0 before (0d), yet not after 4d of LT/EL. Both hexoses, i.e., glucose and fructose, followed similar trends across all genotypes (Fig. 3 C, D). Under control conditions, hexose amounts were highest in *pgm1* and *pc* mutants while, under LT/EL, amounts rose highest in *chup1* and *hc*.

**Figure 3.**
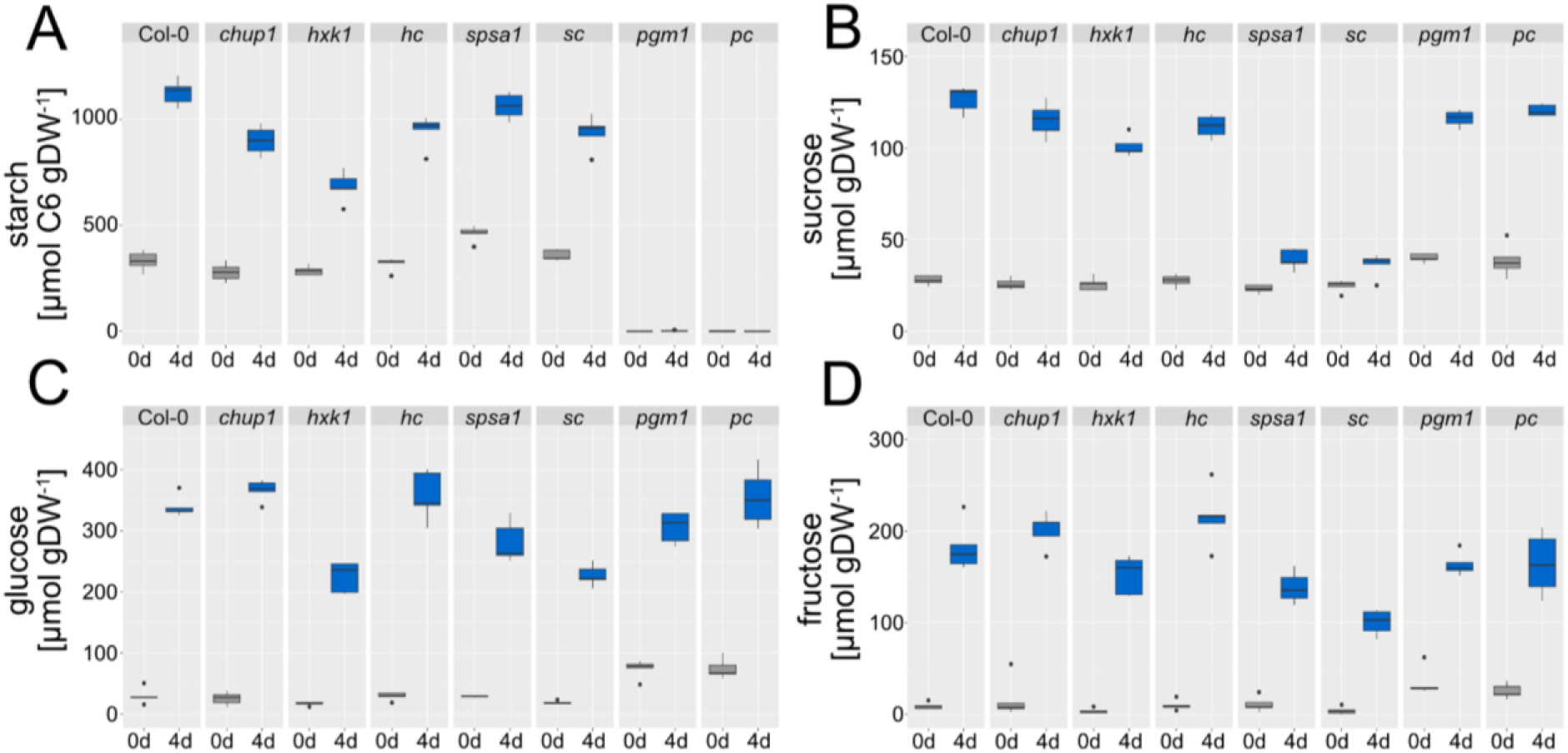
Carbohydrate dynamics under low temperature and elevated light. **(A)** Starch amounts in C6 equivalents. Treatment of 4d LT/EL resulted in a significant increase across all genotypes except for *pgm1* and *pc* (ANOVA, p < 0.001). **(B)** Sucrose amounts before (0d) and after (4d) treatment. Amounts significantly increased in all genotypes due to LT/EL (ANOVA, p < 0.01). **(C)** Glucose amounts before (0d) and after (4d) treatment. Amounts significantly increased in all genotypes due to LT/EL (ANOVA, p < 0.001). **(D)** Fructose amounts before (0d) and after (4d) treatment. Amounts significantly increased in all genotypes due to LT/EL (ANOVA, p < 0.001). n = 5. Grey boxes: control, i.e., 0d of LT/EL. Blue boxes: treatment, i.e., 4d of LT/EL. The full data set is provided in the supplements (Supplementary Table ST1).

In summary, these observations showed that, after 4d at LT/EL, seedlings of *hxk1* mutants were significantly affected in their (above-ground) phenotype, which was also reflected on the molecular level (e.g., anthocyanins, starch and sucrose amounts). Interestingly, this effect was suppressed both on a phenotypic and molecular level in the double mutant *hc*, i.e., when, in addition to HXK1, CHUP1 was also deficient. This suppression was neither observed in *sc* nor in *pc* double mutants, indicating a specific effect between CHUP1 and HXK1 rather than a general effect of (metabolic) acclimation.

### Cytosolic hexoses phosphate amounts increase in *hxk1* under LT/EL

Hexokinase 1 (HXK1) catalyses the phosphorylation of glucose to G6P in a reaction located in the cytosol. To evaluate how this reaction was affected due to the HXK1 mutation, hexokinase activity was determined together with other central enzyme activities and the subcellular distribution of G6P and F6P between the cytosol and plastids (Fig. 4).

**Figure 4.**
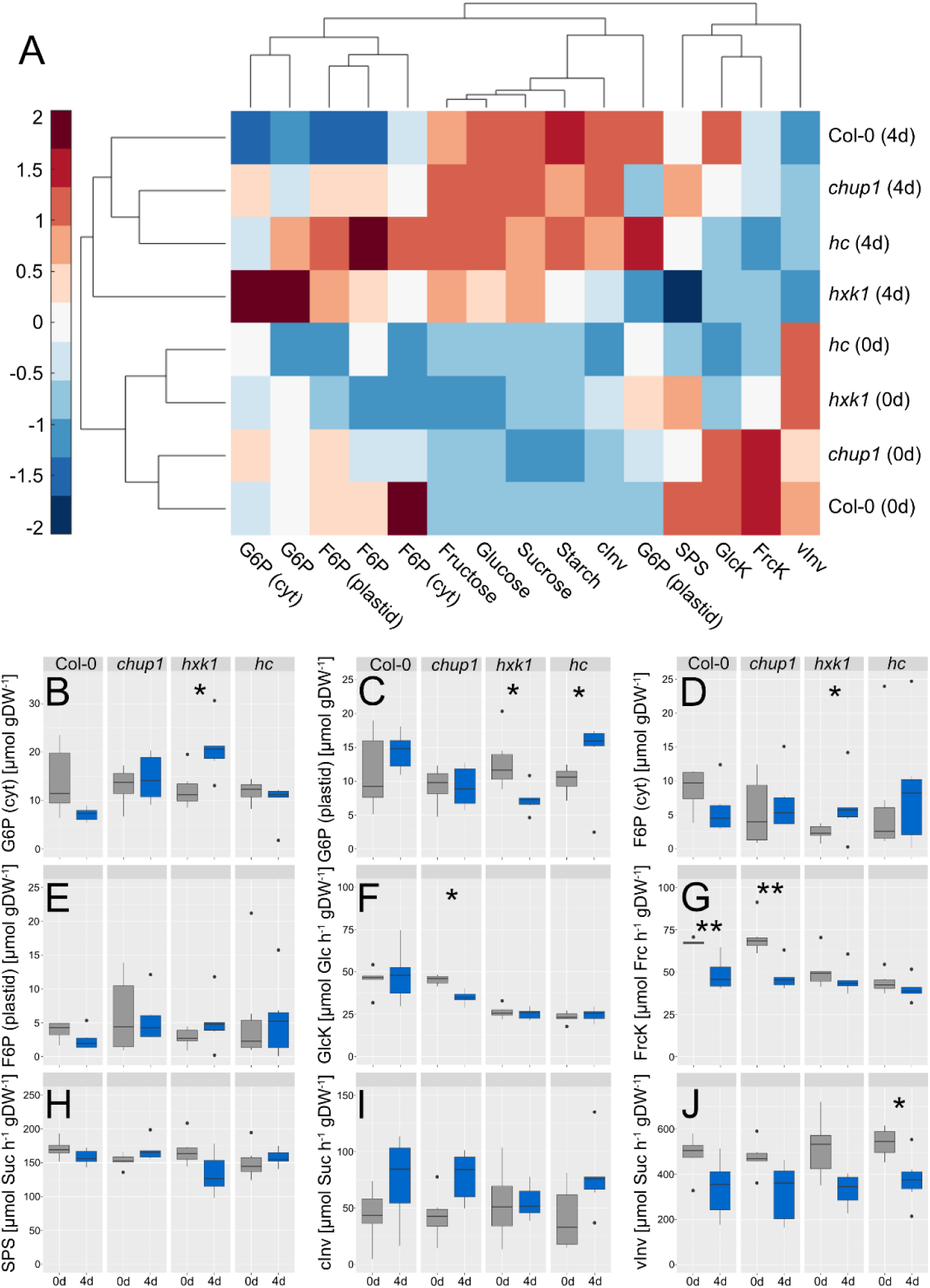
Regulation of carbohydrate metabolism under LT/EL treatment. **(A)** Euclidean distance-based hierarchical clustering of the central carbohydrate metabolism between Col-0, *chup1, hxk1* and *hc*. Trees of samples (rows) and variables (columns) were derived from Euclidean distances of medians for each variable (n = 5-6). Medians were z-scaled (i.e., zero mean/unit variance), and scaled values are presented by the colorbar. **(B)** Amounts of cytosolic G6P, **(C)** Amounts of plastidial G6P, **(D)** Amounts of cytosolic F6P, **(E)** Amounts of plastidial F6P, **(F)** Activity (v_max_) of GlcK, **(G)** Activity (v_max_) of FrcK, **(H)** Activity (v_max_) of SPS, **(I)** Activity (v_max_) of cInv, **(J)** Activity (v_max_) of vInv. Grey boxes: control conditions (0d), blue boxes: 4d LT/EL. Asterisks indicate significance (* p < 0.05, ** p < 0.01, ANOVA/Tukey HSD posthoc test). G6P: glucose 6-phosphate; F6P: fructose 6-phosphate; cInv: cytosolic invertase; vInv: vacuolar invertase; SPS: sucrose phosphate synthase; GlcK: glucokinase; FrcK: fructokinase; cyt: cytosolic. The full data set is provided in the supplements (Supplementary Table ST1).

Under control conditions (0d), carbohydrate metabolism in Col-0 and *chup1* built one cluster while *hxk1* clustered together with *hc*. This was predominantly due to significantly higher FrcK and GlcK activities in Col-0 and *chup1* (ANOVA, p < 0.05), whereas vInv activities were higher in *hxk1* and *hc*. After 4d of LT/EL, *chup1* and *hc* clustered together, while *hxk1* differed most from all other genotypes. The total G6P amount and its cytosolic fraction were most elevated in *hxk1* under these conditions while the plastidial G6P fraction was lower than in Col-0, *chup1* and *hc*. Also, SPS activity was found to be lower in *hxk1* than in the other genotypes.

The most significant effects, which separated *hxk1* from all other genotypes under LT/EL, were (i) increased amounts of cytosolic G6P and F6P, and (ii) decreased G6P amounts in the plastid (Fig. 4 B, C, D). In the double mutant *hc*, cytosolic amounts of G6P and F6P did not differ significantly between 0d and 4d, which reflected the dynamics in Col-0 and *chup1* (Fig. 4 B, D). Plastidial G6P significantly increased in *hc* under LT/EL (Fig. 4 C).

Chlorophyll fluorescence analyses were conducted to evaluate if the observed differences in carbohydrate metabolism were potentially due to, or related to, differential photosynthetic efficiencies. Analyses were only performed on obviously green leaf tissue, i.e., pale/faded leaf area in *hxk1* under LT/EL was excluded from the analysis. Maximum quantum efficiencies of Photosystem II (Fv/Fm) were determined from dark adapted leaves and were found to significantly decrease due to 4d LT/EL in all genotypes (Fig. 5 A). At 4d LT/EL, *hxk1* had a slightly, but significant, lower Fv/Fm than Col-0.

**Figure 5.**
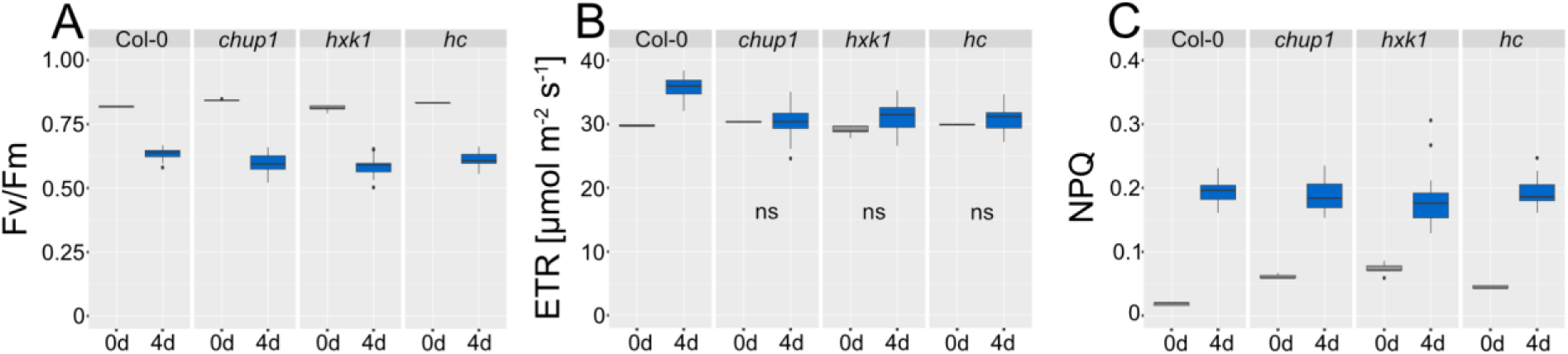
Chlorophyll fluorescence analysis before and after LT/EL treatment. **(A)** Maximum quantum yield of Photosystem II (Fv/Fm); **(B)** linear electron transport rates (ETR); **(C)** non-photochemical quenching (NPQ). Parameters were determined at 22°C and growth PAR intensity, i.e., at 100 μmol photons m^-2^ s^-1^ for control plants and at 250 μmol photons m^-2^ s^-1^ for LT/EL plants. Values of control (0d) and treatment (4d) differed significantly (ANOVA, p < 0.001) within each genotype except for comparisons labelled with ‘ns’ (not significant). Grey boxes: control (0d), blue boxes: LT/EL treatment (4d). n = 5. An overview of all data is provided in the supplements (Supplementary Table ST1).

Comparison of linear electron transport rates (ETR), which were determined after exposure of leaf tissue to growth light intensity, revealed a significant treatment effect only in Col-0 (ANOVA, p < 0.001, Fig. 5 B). Non-photochemical quenching (NPQ) significantly increased due to treatment and reached similar values in all genotypes after 4d at LT/EL (ANOVA, p < 0.001, Fig. 5 C). Interestingly, under control conditions, all mutants had significantly higher NPQ values than Col-0, and *hxk1* had the highest median value of ∼0.75 (Fig. 5 C).

### Deficiency of HXK1 induces an immune response of the transcriptome

Transcript levels were quantified to evaluate how deficiencies of CHUP1, HXK1, or both affect leaf tissue response to LT/EL. In total, ∼32,500 transcribed genes were detected in at least one sample (Supplementary Table ST2). To reveal the most significant effects, transcripts were selected which significantly differed between at least two genotypes or control and treatment, respectively (p < 0.001, ANOVA and TukeyHSD posthoc test incl. correction for multiple testing). This resulted in the identification of 10,094 candidate genes (Supplementary Table ST3). A hierarchical cluster analysis of these significantly affected candidates revealed a group of 1,742 candidate genes of which transcripts were significantly upregulated only in *hxk1* mutants after 4d LT/EL (Fig. 6, black box). Another cluster of 441 candidate genes was detected which were constitutively upregulated in *hxk1* compared to all other genotypes (Fig. 6, blue box). Lists of candidate genes in both clusters are provided in the supplement (Supplementary Table ST 4).

**Figure 6.**
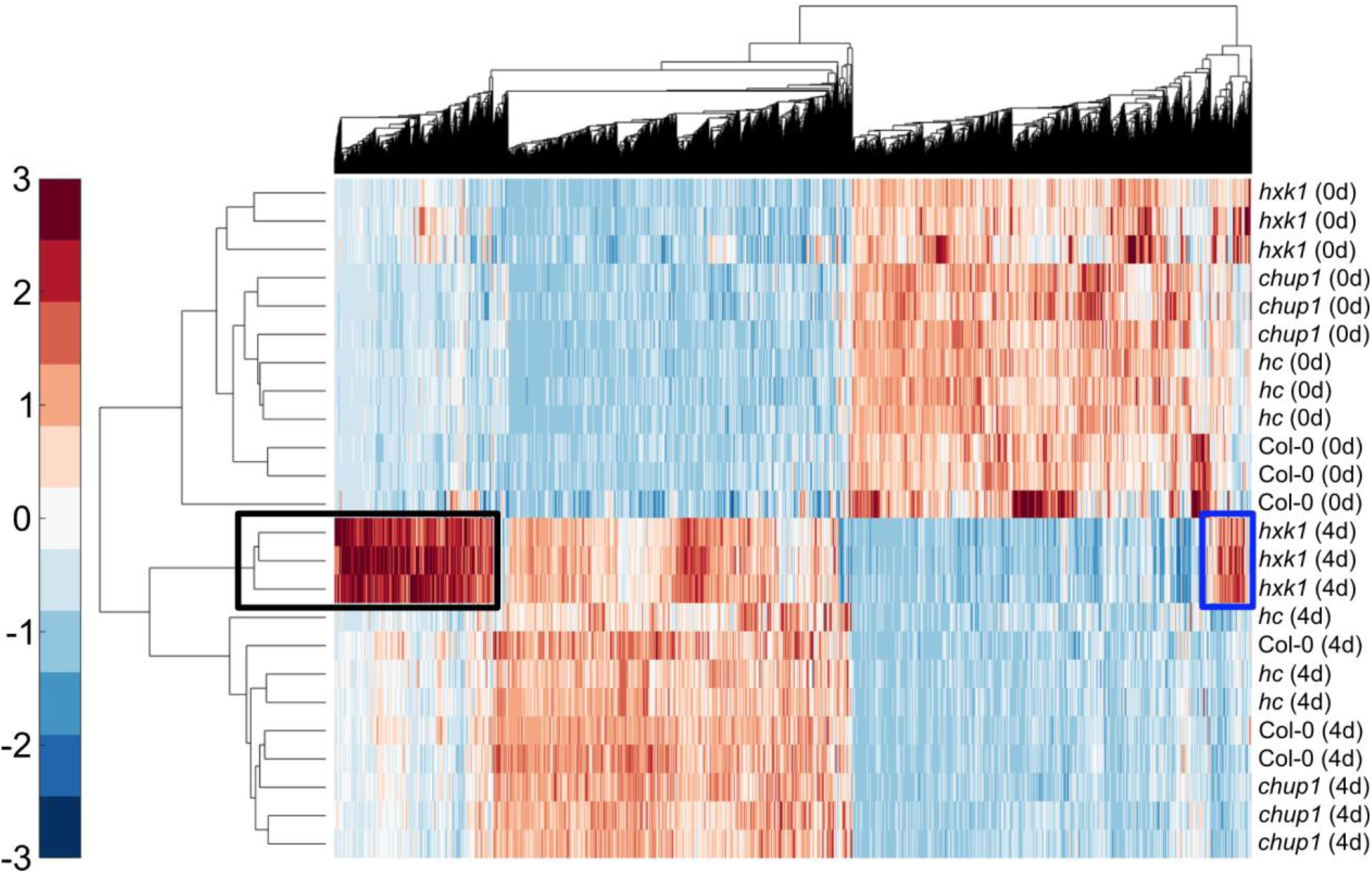
Hierarchical clustering of transcripts and samples before and after 4d at LT/EL. Only transcripts were considered which were, at least, significantly different between conditions or two genotypes (ANOVA, p< 0.001). Each row represents a sample (labels on the right side). Each column represents a candidate gene. Trees were derived from Euclidean distances. The colorbar indicates z-scaled normalised expression values (i.e., zero mean – unit variance scaling). Black box: candidate genes which were only upregulated under LT/EL in hxk1. Blue box: constitutively upregulated candidate genes in *hxk1* under LT/EL. Lists of candidate genes are provided in the supplements (Supplementary Tables ST3 and ST4).

As expected, both *HXK1* and *CHUP1* were among the significantly affected transcripts (Supplementary Figure SF2 A, B). The transcript levels of *HXK1* did not vary significantly in *hxk1* under LT/EL while *CHUP1* transcript levels significantly decreased due to treatment, also in the *chup1* and *hc* mutants. Transcript levels of *PGM1*, which catalyses a central step in starch biosynthesis, strongly and significantly decreased in *hxk1* under LT/EL (Supplementary Figure SF2 D). While transcripts amounts of chalcone synthase, which catalyses the first committed step of flavonoid/anthocyanin biosynthesis, significantly increased in all genotypes under LT/EL, the transcript levels in *hxk1* only increased to ∼30% of the other genotypes (Supplementary Figure SF2 F).

A Gene Ontology (GO) term enrichment analysis showed that genes, which were either constitutively upregulated (Fig. 7 A) or upregulated under LT/EL (Fig. 7 B) in *hxk1*, belonged to the significantly enriched terms of ‘stress response’, defense response’, as well as response to ‘stimulus’, ‘biotic stimulus’ or ‘wounding’.

**Figure 7.**
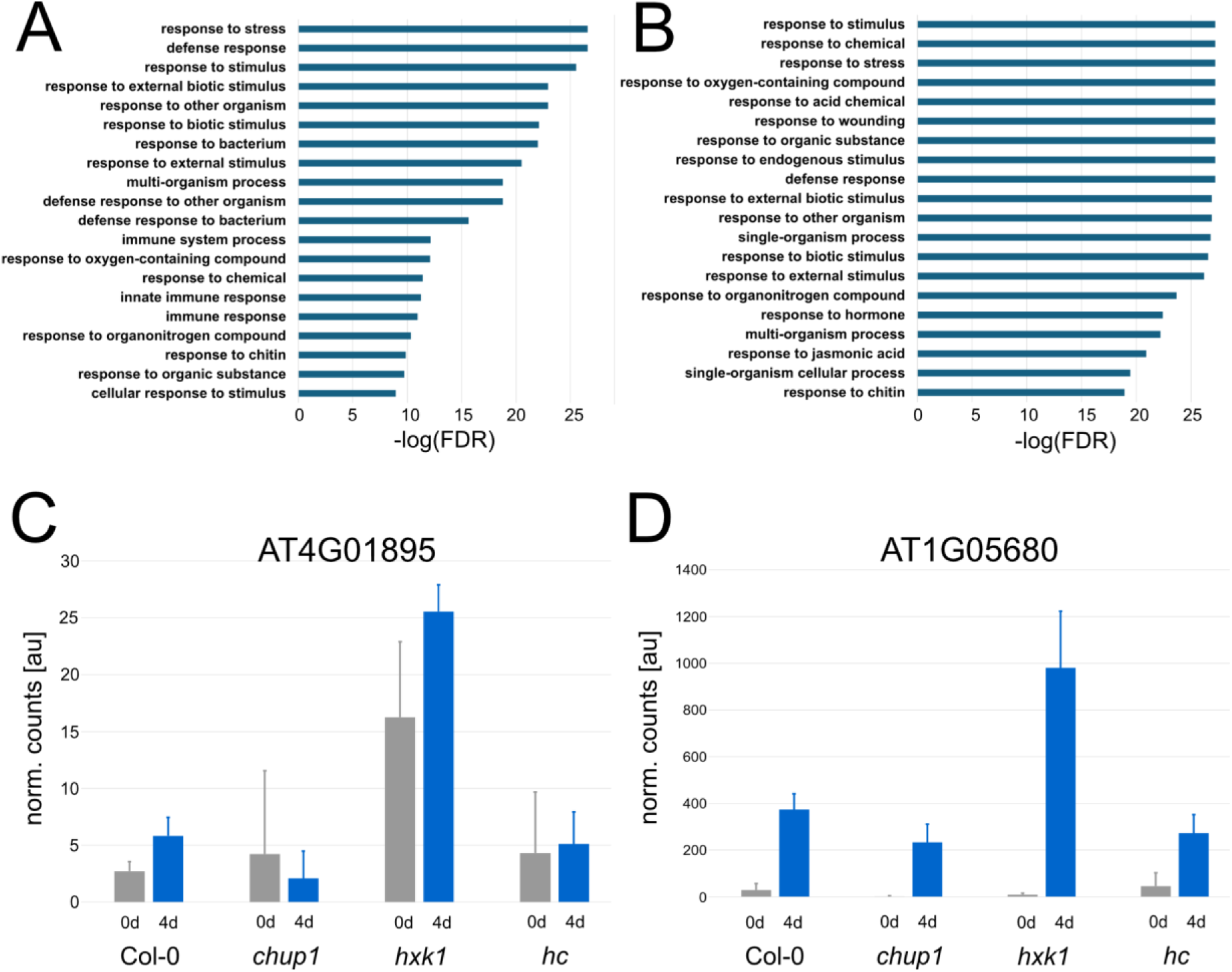
GO term enrichment analysis of upregulated genes in *hxk1*. **(A)** List of the 20 most significantly enriched GO terms among constitutively upregulated transcripts in *hxk1* (corresponds to the blue box in Figure 6; Supplementary Table ST4). **(B)** List of the 20 most significantly enriched GO terms among upregulated transcripts in *hxk1* under LT/EL (corresponds to black box in Figure 6; Supplementary Table ST4). Full lists of significantly enriched terms are provided in the supplements (Supplementary Tables ST5, 6). **(C)** Transcript levels of the *Systemic Acquired Resistance (SAR) regulator protein NIMIN-1-like protein* (AT4G01895), before (0d) and after (4d) LT/EL. **(D)** Transcript levels of *Uridine Diphosphate Glycosyltransferase 74E2* (AT1G05680), before (0d) and after (4d) LT/EL. Bars represent means +/-SD, n = 3.

Further significantly enriched terms were ‘innate immune response’, ‘immune response’, and ‘response to jasmonic acid’. Two proteins involved in biotic stress response whose genes were found to be differentially expressed in *hxk1* due to both constitutive and stress induced dynamics were further analysed (Fig. 7 C, D). Transcript levels of the *Systemic Acquired Resistance (SAR) regulator protein NIMIN-1-like protein* (AT4G01895) were constitutively higher in *hxk1* than in other genotypes (Fig. 7 C). Additionally, transcript levels of *Uridine Diphosphate Glycosyltransferase 74E2* (AT1G05680) were induced by LT/EL across all genotypes, but the response effect was strongest in *hxk1* (Fig. 7 D). In summary, together with the full enrichment analysis, these observations indicated that HXK1 is involved in regulation of transcriptional regulation of plant immune response, and that this response depends on CHUP1, particularly under abiotic stress.

## Discussion

### Carbohydrate metabolism is highly plastic under combined stress of LT/EL

Plant responses to dynamic environmental conditions are multifaceted and depend on a tightly regulated molecular network which coordinates environmental perception with metabolic and signalling responses (Ding et al., 2020). Stabilizing photosynthetic CO_2_ uptake under stressful conditions is of utmost importance for plants to be able to acclimate, grow and propagate in a changing environment. Photosynthetic efficiency and CO_2_ assimilation rates depend, among others, on positioning of chloroplasts and regulation of carbohydrate metabolism (Gotoh et al., 2018; Kitashova et al., 2021; Seydel et al., 2022). Particularly, under stress combinations like low temperature and high light, plants need to efficiently regulate photosynthetic activity with carbohydrate biosynthesis to prevent limitations in electron transport, ATP biosynthesis or redox metabolism (Geigenberger et al., 2017; Kleine et al., 2021; Schwenkert et al., 2022). In the present study, both cell biological and metabolic functions and pathways were mutated to reveal their roles in stabilising photosynthesis in a LT/EL regime. A deficiency of SPS activity resulted in a strongly affected sucrose accumulation response under stress conditions in both *spsa1* and *sc* mutants. This suggests that affected chloroplast positioning due to CHUP1 deficiency does not amplify nor release the limitation of sucrose biosynthesis in a *spsa1* background. This indicates that metabolite transport across the chloroplast envelope, e.g., catalysed by TPT for triose phosphate supply in the cytosol, seems not to be permanently affected by the spatially dense accumulation of chloroplasts (see Supplementary Fig. SF1). A mutation of *PGM1* resulted in significantly elevated sugar amounts under control conditions, which indicates the shift of carbon partitioning from starch biosynthesis to sucrose biosynthesis and other anabolic pathways (Gibon et al., 2009; Brauner et al., 2014). However, as already observed for *spsa1*, additional CHUP1-deficiency did not significantly affect the carbohydrate amounts of *pgm1*, neither under control nor under stress conditions. Hence, in addition to the cytosolic sucrose biosynthesis pathway, the chloroplast-located pathways of carbohydrate metabolism also did not seem to be significantly affected under analysed conditions by the polar chloroplast accumulation in the cells. In summary, these findings indicate a high plasticity of photosynthesis and carbohydrate metabolism under LT/EL that is stabilized even with affected localization of photosynthetic organelles, which has been shown to have significant impact on biomass accumulation (Gotoh et al., 2018).

### Exposure to combined LT/EL increases susceptibility of the *hxk1* mutant for biotic stress

In contrast to *spsa1* and *pgm1* mutants, the *hxk1* mutant did not accumulate (photometrically measurable) anthocyanins under LT/EL (see Fig. 2 D). Further, under LT/EL, the *hxk1* mutant had a lower ratio of fresh weight to dry weight, significantly lowered chlorophyll content and less starch, sucrose and glucose than Col-0. Together with the growth phenotype, which showed lesions and/or pale leaf areas, this suggested a significant impact on photosynthesis and carbohydrate metabolism. The quantification of subcellular hexose phosphates revealed an overaccumulation of G6P in the cytosol of *hxk1* while the plastidial fraction was depleted under LT/EL. Together with the significantly reduced *PGM1* and *BAM3* transcripts, this suggests that reduced starch amounts in *hxk1* resulted rather from a combination of substrate depletion and downregulation of the starch biosynthesis pathway than an increased rate of starch degradation.

Surprisingly, chlorophyll fluorescence analysis of the (obviously) green leaf areas revealed only a slight, yet still significant, reduction of Fv/Fm in *hxk1* compared to Col-0. The significance of this effect was reduced in *hc* double mutants (i.e., p > 0.05), suggesting a stabilising effect of the *CHUP1* mutation in the *hxk1* background. In contrast, while Col-0 significantly increased ETR after 4d LT/EL, the ETRs of treated *chup1*, *hxk1* and *hc* mutants did not differ from those of the control. This missing acclimation of photosynthetic electron transport might have different causes: in *chup1* a missing relocation of chloroplasts probably limits the efficiency of photochemistry (Gotoh et al., 2018), while in *hxk1* it might be that the observed reduction of carbon partitioning into starch and other metabolic pathways feedback inhibits and/or limits the rate of electron transport. Although overexpressing, and not attenuating, HXK1 has been found to inhibit photosynthesis (Dai et al., 1999; Granot et al., 2014), it might be that due to the combined LT/EL stress condition, other effects or regulatory factors caused a limitation or repression of photosynthetic acclimation. In this context, the analysis of the transcriptome revealed a strong and highly significant enrichment of transcripts which belong to biotic stress and immune response categories in *hxk1* under LT/EL. Based on findings which suggested HXK1 to be a positive regulator of plant immunity (Jing et al., 2020), this indicates that *hxk1* is susceptible to fungal or bacterial infection which caused the strong transcriptional response. Under control conditions, such a response was already observed, but it was significantly amplified by the 4d LT/EL treatment. This suggests a role of HXK1 in integrating and regulating plant responses towards biotic and abiotic stressors simultaneously. The finding of an absent accumulation of anthocyanins and a reduced expression of *CHS* in *hxk1* (see Fig. 1 and Supplementary Figure SF2 F) further supports this hypothesis due to the previously reported repression of flavonoid accumulation by microbe-associated molecular patterns triggered immunity (Serrano et al., 2012).

The reversible nature of the growth and metabolic phenotype of *hxk1* in the *hc* double mutant suggests an interference of chloroplast positioning, mediated by CHUP1, and HXK1 function. Interestingly, a previous study provided evidence for a role of (epidermal) chloroplast positioning in controlling the entry of fungal pathogens (Irieda and Takano, 2021). The authors showed that blocking this epidermal chloroplast response significantly decreases preinvasive nonhost resistance of *Arabidopsis* against fungi. Based on their experiments, they suggested that CHUP1 is a negative regulator of the epidermal chloroplast response. Thus, a reason for the absence of the *hxk1* phenotype in the *hc* double mutant in the present study may be that the deficient positive immune regulator HXK1 is counteracted by a deficiency of the negative regulator CHUP1 (Fig. 8).

**Figure 8.**
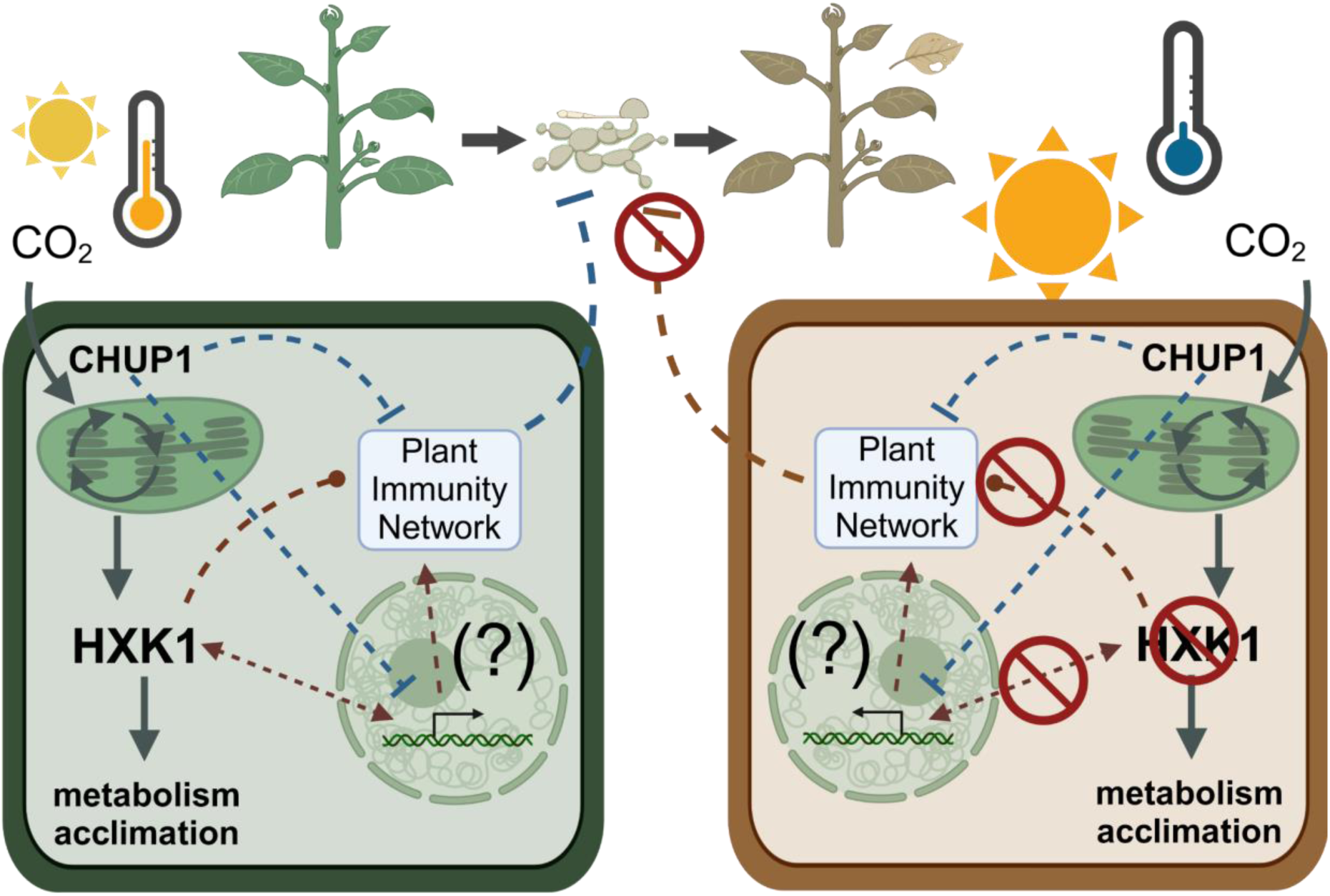
Suggested model to connect HXK1 and CHUP1 to the plant immune network. Solid lines indicate carbon fluxes, dashed lines indicate regulatory interactions. CHUP1 acts as a negative regulator of plant immunity and HXK1 as a positive regulator. Simultaneously, both have a function in photosynthesis, metabolic regulation and acclimation to a changing environment. In *hxk1* mutants, CHUP1 negatively affects plant immunity which makes the plants more susceptible to biotic stress. This is amplified by abiotic stress, e.g., LT/EL. Left side: wild type model, right side: model for *hxk1* plants under LT/EL. Created in https://BioRender.com.

These findings clearly indicate that photosynthesis, the carbohydrate metabolism and the response toward biotic and abiotic stressors are integrated in a regulatory and signalling network in which HXK1 plays a central role. Finally, due to the observation that *chup1* rescued the *hxk1* phenotype, knocking out or down CHUP1 might represent a (conserved) strategy to improve plant immunity which needs to be evaluated in future studies.

## Acknowledgements

We thank the members of Plant Evolutionary Cell Biology at LMU München and the members of TRR175 for constructive discussions and advice. We thank Prof. Andreas Klingl for support with the TEM analysis. Further, we thank the Graduate School Life Science Munich (LSM) for support. This work was funded by Deutsche Forschungsgemeinschaft (DFG), TRR175/C01 and TRR175/D03.

## Conflict of interest statement

The authors declare no conflict of interest.

## Data availability statement

Data of this study is available in the supplements and in a repository of the NFDI Data Plant consortium: <u>Link to Repository</u>

## Author contributions

**SBB** performed subcellular fractionation and established subcellular hexose phosphate quantification assay, performed cellular and subcellular quantification of metabolites, statistics and data evaluation, and wrote the manuscript. **AK** generated plant material, performed enzyme maximal activity quantification, performed statistics and data evaluation, and wrote the manuscript. **BÖ** generated plant material, performed genotyping and enzyme activity quantification. **CKY** performed subcellular fractionation. **BES** performed microscopy. **LS** generated double mutants. **TK** performed transcriptomics analysis. **TN** conceived the study, performed statistics and data evaluation, and wrote the manuscript. All authors reviewed and approved the manuscript.

## Supplementary Information

**Supplementary Figure SF1.**
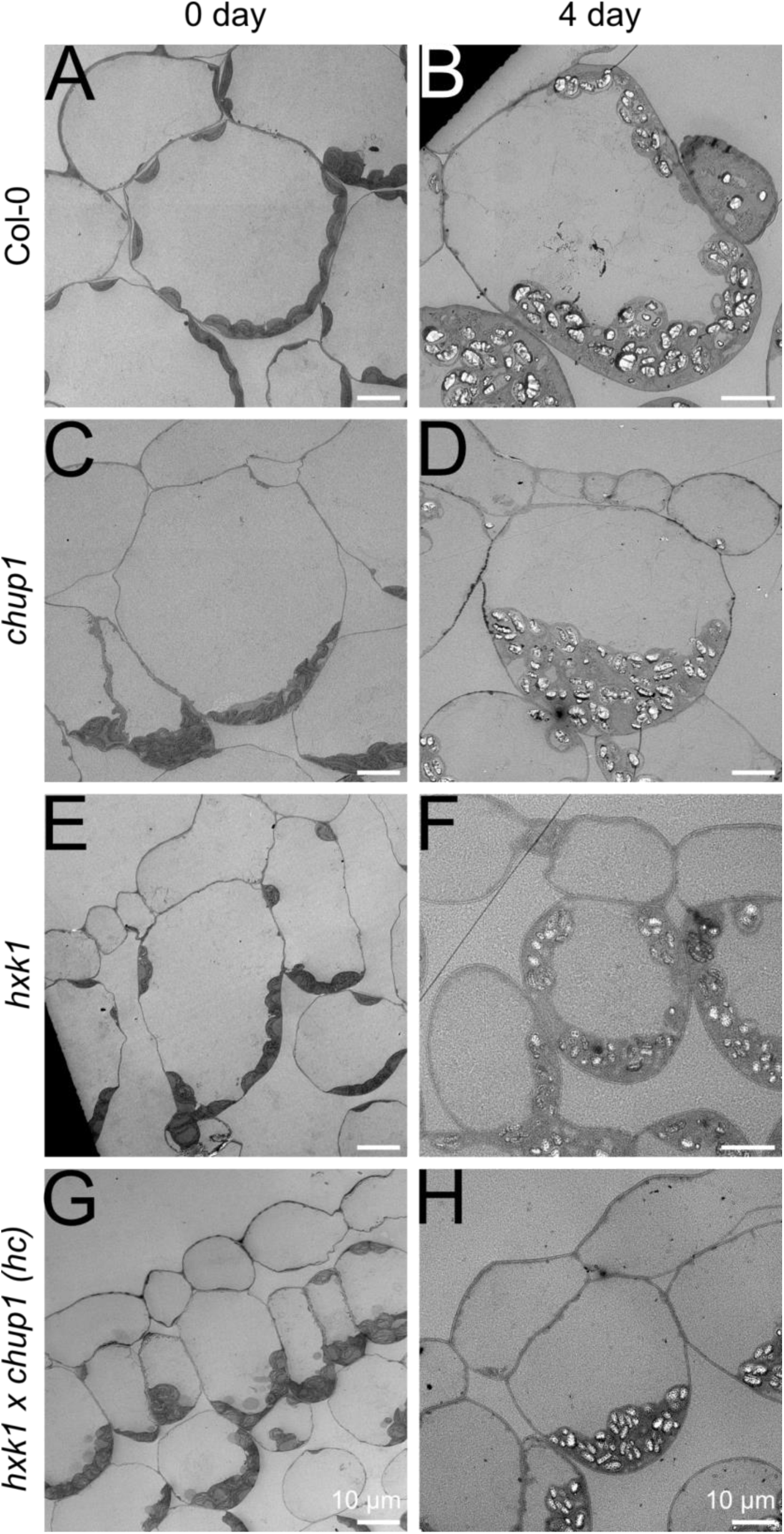
Dynamics of chloroplast positioning. Transmission electron microscopy of leaf tissue in: **(A)** and **(B)** Col-0, **(C)** and **(D)** *chup1*, **(E)** and **(F)** *hxk1*, **(G)** and **(H)** *hc*. The left panel (**A, C, E, G**) shows chloroplast positioning before LT/EL treatment (0d). The right panel (**B, D, F, H**) shows chloroplast positioning after 4d at LT/EL.

**Supplementary Figure SF2.**
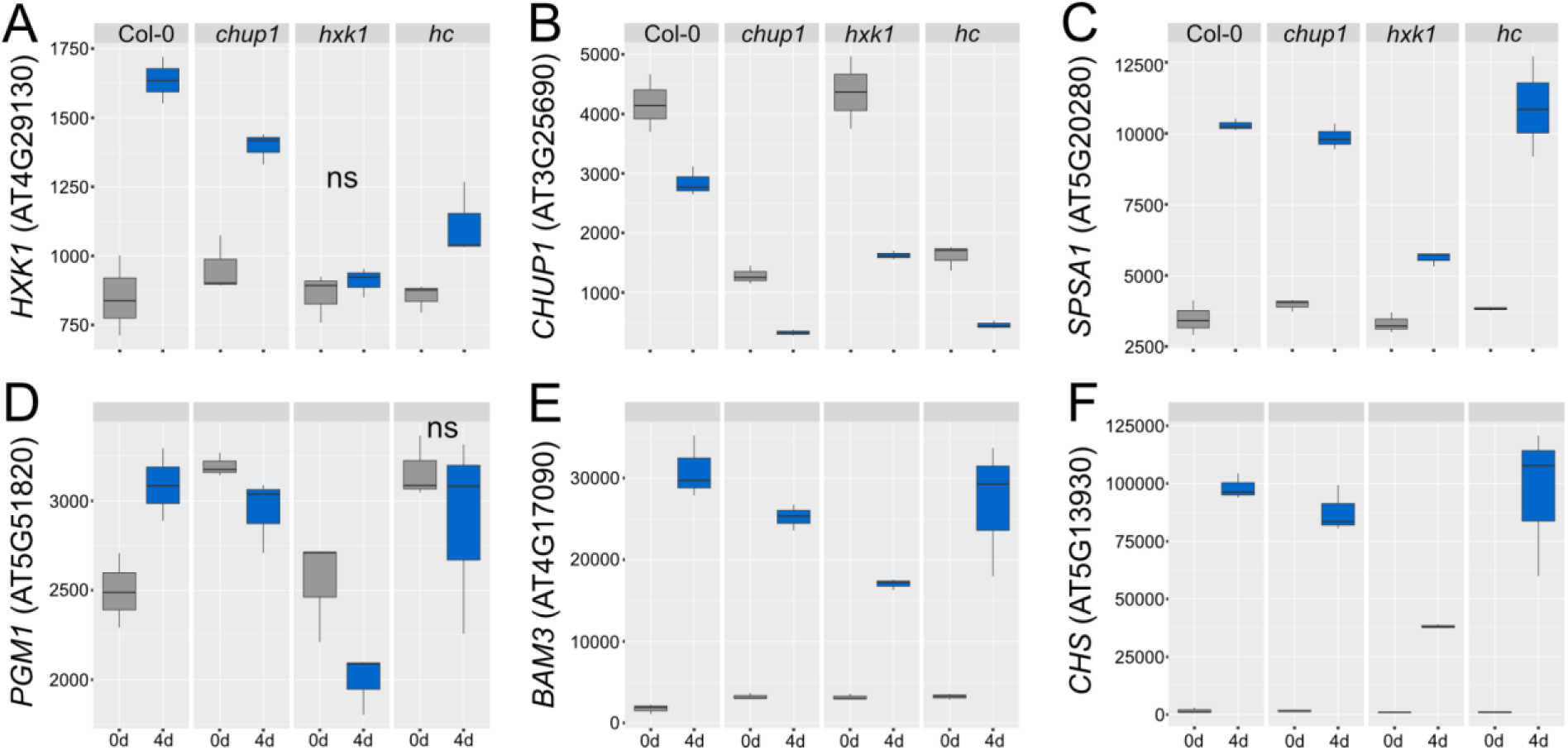
Transcripts of selected candidate genes with a role in chloroplast positioning, primary and secondary metabolism. (A) Hexokinase 1, (B) Chloropalst Unusual Positioning 1, (C) Sucrose Phosphate Synthase A1, (D) Phosphoglucomutase 1, (E) beta-Amylase 3, (F) Chalconsynthase. All differences between 0d and 4d were significant in each genotype except for those labeled with ‘ns’ (p < 0.001, ANOVA). Grey boxes: 0d; blue boxes: 4d. n= 3.

**Supplementary Table ST1. Experimentally determined variables of growth, metabolism and chlorophyll fluorescence.** Chlorophyll content [mg gDW^-1^]; protein content [mg gDW^-1^]; anthocyanins [μmol C15 gDW^-1^]; starch [μmol C6 gDW^-1^]; all other metabolite amounts [μmol gDW^-1^]; all enzyme activities (v_max_) [μmol h^-1^ gDW^-1^].

**Supplementary Table ST2. Full list of transcript levels detected in at least one sample.**

**Supplementary Table ST3. Transcripts which changed significantly at least between two genotypes or between treatments** (p < 0.001, ANOVA and TukeyHSD posthoc test incl. correction for multiple testing).

**Supplementary Table ST4. Candidate genes which were (i) upregulated under LT/EL or (ii) constitutively upregulated in *hxk1*.**

**Supplementary Table ST5. Summary of GO term enrichment analysis of constitutively upregulated candidate genes in *hxk1*.**

**Supplementary Table ST6. Summary of GO term enrichment analysis of LT/EL-induced candidate genes in *hxk1*.**

## Notes

https://git.nfdi4plants.org/thomas.naegele/Bagshaw_et_al_HXK1_CHUP1

### Competing Interest Statement

The authors have declared no competing interest.

